# Evaluating the feasibility and efficacy of a dual-modality nanoparticle contrast agent (Nanotrast-CF800) for image-guided sentinel lymph node mapping in the oral cavity of healthy dogs

**DOI:** 10.1101/2021.07.26.453824

**Authors:** Jennifer Wan, Michelle L. Oblak, Ann S. Ram, Charly McKenna, Ameet Singh, Stephanie Nykamp

## Abstract

A combination of pre- and intraoperative sentinel lymph node (SLN) mapping techniques has been suggested to optimize SLN detection. A novel liposomal nanoparticle, Nanotrast-CF800 (CF800), utilizes computed tomography lymphography (CTL) and near infrared fluorescence imaging (NIRF) for image-guided surgery and SLN mapping. This novel tracer agent has not been evaluated in companion animals. The objective of this study was to evaluate the feasibility and efficacy of CF800 for SLN mapping in the oral cavity of healthy dogs and to report any local adverse effects. Six healthy adult purpose-bred research dogs randomly received either 1 mL (group 1) or 2 mL (group 2) of CF800 injected into the submucosa at the level of the right canine maxillary tooth. CTL and percutaneous NIRF were performed at 1, 3, and 10 minutes, then 1, 2, 4, 7, and 10 days post-injection (p.i). Overall, both CTL and NIRF identified SLNs in all dogs. The overall peak mean contrast enhancement of the SLNs was 73.98 HU (range 63.45-86.27 HU) at 2 days p.i. Peak fluorescence of the SLN occurred at 1 day p.i. The agent was retained within the SLN for at least 7 days for CTL and 4 days for NIRF. No adverse effects were observed. Local administration of CF800 was simple and feasible for the detection of SLNs using CTL+NIRF in the head and neck of healthy dogs and was not associated with significant local adverse events.

## Introduction

Lymph node assessment for metastatic disease is integral for staging many solitary cancers as it provides valuable prognostic information and can guide treatment recommendations. The SLN is defined as the first lymph node(s) that drains a primary tumor and is based on the theory that metastasis occurs in an orderly and sequential manner through the regional lymphatic basin.^1,2^ The SLN has been demonstrated to be predictive of metastatic disease.^3–5^ A negative SLN suggests that there has been no further systemic spread and requires no additional therapy while a SLN positive for metastasis can negatively impact a patient’s prognosis.^1–4^ Biopsy of the SLN is a less invasive surgical technique for local tumor control that reduces patient morbidity.^6,7^

Several mapping techniques have been described to localize and identify the SLN, and historically, included dyes^8,9^ and lymphoscintigraphy.^10^ The combination of lymphoscintigraphy and blue dye is currently the standard of care for SLN mapping in human oncology; however, there are limitations associated with these techniques.^2,11,12^ The use of blue dyes has the potential to cause severe allergic reactions^8,13^, while lymphoscintigraphy requires the use of specialized equipment, is associated with radiation exposure, and has higher costs.^2^

Computed tomography is a rapid and noninvasive diagnostic modality that provides detailed anatomic information and high spatial resolution for accurate preoperative planning. Computed tomography lymphography (CTL) is an alternative SLN mapping modality that utilizes a water soluble based iodinated contrast, which results in the rapid identification of contrast-enhanced lymphatic tracts and SLNs.^14–16^ While this modality has excellent utility for preoperative mapping, it does not provide the surgeon with intraoperative guidance, thus, CTL should be combined with an optical imaging modality to ensure accurate intraoperative SLN identification.^15,17^

There is growing interest in the use of near infrared fluorescence imaging (NIRF) with indocyanine green (ICG) for SLN mapping modality in both human and veterinary medicine. This technique provides real-time optical intraoperative guidance for the detection of fluorescent SLNs. Its use has been favorable due to its ease of application, high SLN detection rates, and association of ICG with reasonable cost and minimal adverse effects.^2,18,19^ In a 2016 metaanalysis review, the use of ICG alone for SLN mapping in human breast cancer patients yielded an overall SLN detection rate of 98% with a high sensitivity and specificity and low false negative rate.^5^ A potential limitation of ICG is associated with its rapid migration through the lymphatic system. ICG has a small molecular weight and binds to plasma proteins, which allows it to become quickly distributed throughout the lymphatic system.^20–22^ As a result, ICG may not retain within the SLN and could migrate to second- and third-tier lymph nodes causing misidentification of the true SLN and leading to the unnecessary excision of additional lymph nodes.^21^ Furthermore, NIRF SLN mapping needs to be performed almost immediately after contrast injection.

A combination of preoperative and intraoperative SLN mapping techniques, such as CTL with intraoperative NIRF, has been suggested to facilitate the accurate detection of SLNs. A limitation of this technique is the need to perform two separate injections. Due to the time that elapses between preoperative imaging and surgery, there can be significant time between these injections and often different operators will perform the injections. As a result, there may be variability in the location and injection techniques which could alter their agreement accuracy. A novel dual-imaging modality tracer agent Nanotrast-CF800 (CF800) was recently developed and evaluated in a preclinical animal model.^23^ This novel tracer incorporates both iohexol and ICG within a liposomal nanoparticle capsule (mole ratio 1000:1) for the use of preoperative and intraoperative image-guided tumor and metastatic lymph node localization. Results of this study demonstrated CT contrast enhancement and fluorescence intensity within metastatic cervical lymph nodes up to 4 days p.i. in rabbits with induced tumorigenesis of the oral cavity. The authors proposed that this novel agent can be administered to utilize both preoperative and intraoperative SLN mapping for staging various cancers while achieving an increased retention time within the targeted tissues and allow for appropriate surgical planning. The use of this novel agent has not yet been investigated in companion animal patients.

The main objective of this experimental study is to evaluate the feasibility and efficacy of dual-modality nanoparticle CF800 for image-guided SLN mapping in the oral cavity of healthy dogs. A secondary objective is to report any local adverse effects associated with CF800. We hypothesized that the use of CF800 will be feasible and effective for SLN mapping within the oral cavity of healthy dogs and the local administration within the oral cavity will not result in local adverse events.

## Materials and Methods

### Animals

Six adult purpose-bred research beagle dogs were utilized for this experimental study. This research project was approved by the University of Guelph Animal Care Committee (AUP ~3775). All dogs were determined to be healthy based on physical examinations, complete blood count, serum biochemical profile, and urinalysis prior to the start of the study. Dogs were randomly divided into two groups to receive either 1 mL (group 1) or 2 mL (group 2) of CF800.

### Study design

Dogs were fasted for at least 8 hours prior to the start of the study. On the day of the procedure, dogs were sedated using 0.05 mg/kg hydromorphone and 5-10 mcg/kg dexmedetomidine IV via the cephalic or lateral saphenous vein. Dogs were positioned in left lateral recumbency on the CT scanner table and the fur from the ventral mandible to thoracic inlet was clipped. Cardiorespiratory parameters including heart rate, respiratory rate, temperature, and indirect oscillometric blood pressures were monitored from starting at 10-15 minutes prior to injection then every 5 minutes following injection until recovery from sedation. All dogs underwent a pre-contrast CT scan extending from the nose to the thoracic inlet using a 16-slice detector CT (GE Brightspeed CT scanner, GE Healthcare, Milwaukee, Wisconsin, United States) with data collected using 0.625mm slice thickness and standardized protocol in helical mode, 0.8 second rotation time, collimator pitch of 1, 120 kV and 200mAs. A near infrared (NIR) exoscope (VITOMII, Karl Storz Endoscopy Canada Ltd., Mississauga, ON, Canada) was positioned 20 cm above the patient’s oral cavity and ventral neck and connected to a light source and display monitor.^24^ Based on the assigned group, CF800 was injected using a 25g needle into the labial mucosa at the dorsal aspect of the right canine maxillary tooth over 1 minute, as previously described by Townsend et al. (2019).^25^ Fluorescence was observed continuously during injection. Following injection, CT scans of the head and neck were performed at 1, 3, and 10 minutes p.i. using the previously described parameters. In between CT scans, NIRF was observed to evaluate the injection site and ventral cervical region for the presence of percutaneous fluorescent lymphatic tracts and right mandibular lymph nodes. The SLN was defined as the first lymph node that was contrast-enhancing on CT and/or the presence of percutaneous fluorescence. Second- and third-tier SLNs were defined as lymph nodes that were not contrast enhanced when the SLN(s) was identified but were enhanced on subsequent scans. The degree of fluorescence intensity was determined based on a semi-quantitative scoring system on a scale from 0 − 3+ (0 = no fluorescence, 1+ = mild fluorescence, 2+ = moderate fluorescence, 3+ = marked fluorescence) ^26^ Videos and images were recorded from the NIR exoscope. Dogs were then reversed using atipamezole IM.

For all subsequent imaging, dogs were sedated using the same protocol as the day of the procedure. Repeat CT scans and NIRF of the head and neck were performed on days 1, 2, 4, 7, and 10 p.i. Daily physical examination and assessment and documentation of the local injection site were performed. Repeat complete blood count and serum biochemistry profiles were performed on day 7. Dogs were returned to the research colony at the conclusion of the study.

### Outcome measures

#### CT lymphography

Images from the CT scans were reviewed and analyzed using Horos Imaging Software (v1.1.7, Open Source Licence; Version 3 (LPGL-3.0). Images of the injection site were evaluated in a transverse plane and bone window (WW: 2000; WL: 350) while images of the right cervical lymph nodes (mandibular and medial retropharyngeal) were evaluated in a multiplanar and soft tissue window (WW: 400; WL: 40) reconstruction. The number and location of the right mandibular lymph nodes was recorded. The SLN and time to identification were recorded.

The injection site was assessed for the following measures:

1. Attenuation of contrast enhancement measured in Hounsfield units (HU) in pre- and post-injection studies at each time point. This was determined by placing a representative region of interest (ROI) within a maximally enhancing region of the injection site and the mean HU was recorded. Three measurements were performed for each dog at each time point and averaged.
2. The ROI volume of contrast was determined by manually drawing the borders of contrast enhancement every 2-3 slices moving from rostral to caudal through the imaging studies at each time point. The mean ROI volume was recorded.

The right cervical lymph nodes (mandibular and medial retropharyngeal nodes) were assessed for the following measures:

1. Attenuation of contrast enhancement measured in HU in pre- and post-injection studies at each time point. A representative ROI was placed at a maximally enhancing region within the lymph node and the mean HU was recorded.
2. The ROI volume of contrast within the lymph node was determined by manually drawing the outline of each lymph node every 2-3 slices moving from rostral to caudal through the imaging studies at each time point. The mean ROI volume of each lymph node was recorded.
3. The length and height of each lymph node was measured in the sagittal and dorsal planes at its largest cross-sectional area, respectively. The width of each lymph node was measured in the transverse plane at its largest cross-sectional area. These measurements were used to calculate the volume of each lymph node using the following formula for calculating the volume of an ellipsoid:

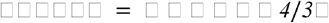 The lymph node volume of each lymph node at each time point was recorded.

#### NIRF

Images captured from the NIR exoscope were exported to a workstation (Microsoft Surface, Microsoft Corporation, Redmond, WA, USA) and reviewed using an open source imaging program (ImageJ). Data collected include the presence of percutaneous fluorescence of the right mandibular lymphocentrum, time to detection of the SLN, the presence of percutaneous fluorescent lymphatic tracts, and the degree of fluorescence of the right mandibular lymphocentrum. The fluorescence intensity at the injection site was measured by outlining the borders of fluorescence and the corrected total fluorescence was measured in pixels.^26^

### Statistical analysis

Descriptive statistics were used to determine the time to detection of SLNs, the number of SLNs on CTL, contrast enhancement of the injection site and SLNs, degree of fluorescence intensity at the injection site and right mandibular lymph node, ROI volume of the injection site and cervical lymph nodes, and lymph node volume of the cervical lymph nodes. Categorical variables were measured as proportions (i.e. presence of percutaneous lymphatic tracts).

For statistical analysis of CT images, the right cervical lymph nodes were grouped as a single unit (if there were ≥1 SLNs identified) and defined as an overall SLN. The mean peak HU and time to peak HU was identified for the injection site and SLNs. Analysis of Variance (ANOVA) of repeated measures was performed to compare mean contrast enhancement and mean ROI volume at the injection site and SLN between groups and over time. The mean lymph node volume was also compared between groups and over time. ANOVA was also used to determine the mean peak HU at the injection site and SLNs and median peak fluorescence of the right mandibular lymph node and compared between groups.

A general linear mixed model for binary distribution was used to compare the degree of fluorescence intensity at the injection site and SLN between groups and over time.

## Results

### CTL

A summary of findings from CTL is presented in Table 1. Sentinel lymph nodes were identified in all dogs and were detected at a median of 5.5 minutes (range 1-1440 minutes). All SLNs identified were ipsilateral to the site of injection. There was a median of 2 SLNs (range 1-3) identified. The right dorsolateral mandibular lymph node (DLM) was identified as sentinel in all dogs (Figure 1). In 2 dogs, the right ventromedial mandibular lymph node (VMM) was also identified, and in another 2 dogs, both the right VMM lymph node and medial retropharyngeal lymph nodes (MRP) were also sentinel. One dog from group 2 had a parotid lymph node identified as sentinel in addition to the DLM. Second-tier SLNs were identified in 4/6 dogs at a median of 24 hours p.i. The VMM node was identified in 2 dogs and the MRP in all four dogs. A third-tier lymph node was identified in only 1 dog, which was the MRP. Contrast attenuation could be identified within the right DLM at 7 days p.i. in all dogs. By 10 days p.i. 5/6 dogs had contrast visible within the right DLM. Contrast-enhanced lymphatic tracts were identified in 2/6 dogs and were only identified in group 2 dogs at 1 minute p.i. (Figure 2).

**Table 1.** Summary of general CTL and NIRF results

|  | CTL |  |  |  |  |  | NIRF |  |  |  |
| --- | --- | --- | --- | --- | --- | --- | --- | --- | --- | --- |
| Dog | SLN Identified (Y/N) | Time to SLN Identification | SLN Identified | Mean ROI Injection (cm<sup>3</sup>) | Mean ROI LN (cm<sup>3</sup>) | Mean LN Volume (cm<sup>3</sup>) | SLN Identified (Y/N) | Time to SLN Identification | *Degree of Fluorescence | Presence of Lymphatic Tracts (Y/N) |
| Group 1 |  |  |  |  |  |  |  |  |  |  |
| 1 | Y | 1 min | R DL mandibular | 0.94 | 0.27 | 2.98 | Y | 8 min | 2+ | N |
| 2 | Y | 10 min | R DL mandibular |  |  |  | Y | 24 h | 2+ | N |
| 3 | Y | 1 min | R DL & VM mandibular, R MRP |  |  |  | Y | 5 min | 1+ | N |
| Group 2 |  |  |  |  |  |  |  |  |  |  |
| 4 | Y | 10 min | R DL & VM mandibular | 1.20 | 0.18 | 2.14 | Y | 9 min | 1+ | Y |
| 5 | Y | 1 min | R DL & VM mandibular |  |  |  | Y | 5 min | 3+ | Y |
| 6 | Y | 24 h | R DL & VM mandibular, R MRP |  |  |  | Y | 6 min | 2+ | Y |

Abbreviations: DL = dorsolateral; VM = ventromedial; MRP = medial retropharyngeal; LN = lymph node; ROI = region-of-interest; SLN = sentinel lymph node
\*Degree of fluorescence: 0 = none; 1+ = mild; 2+ = moderate; 3+ = marked

**Figure 1.**
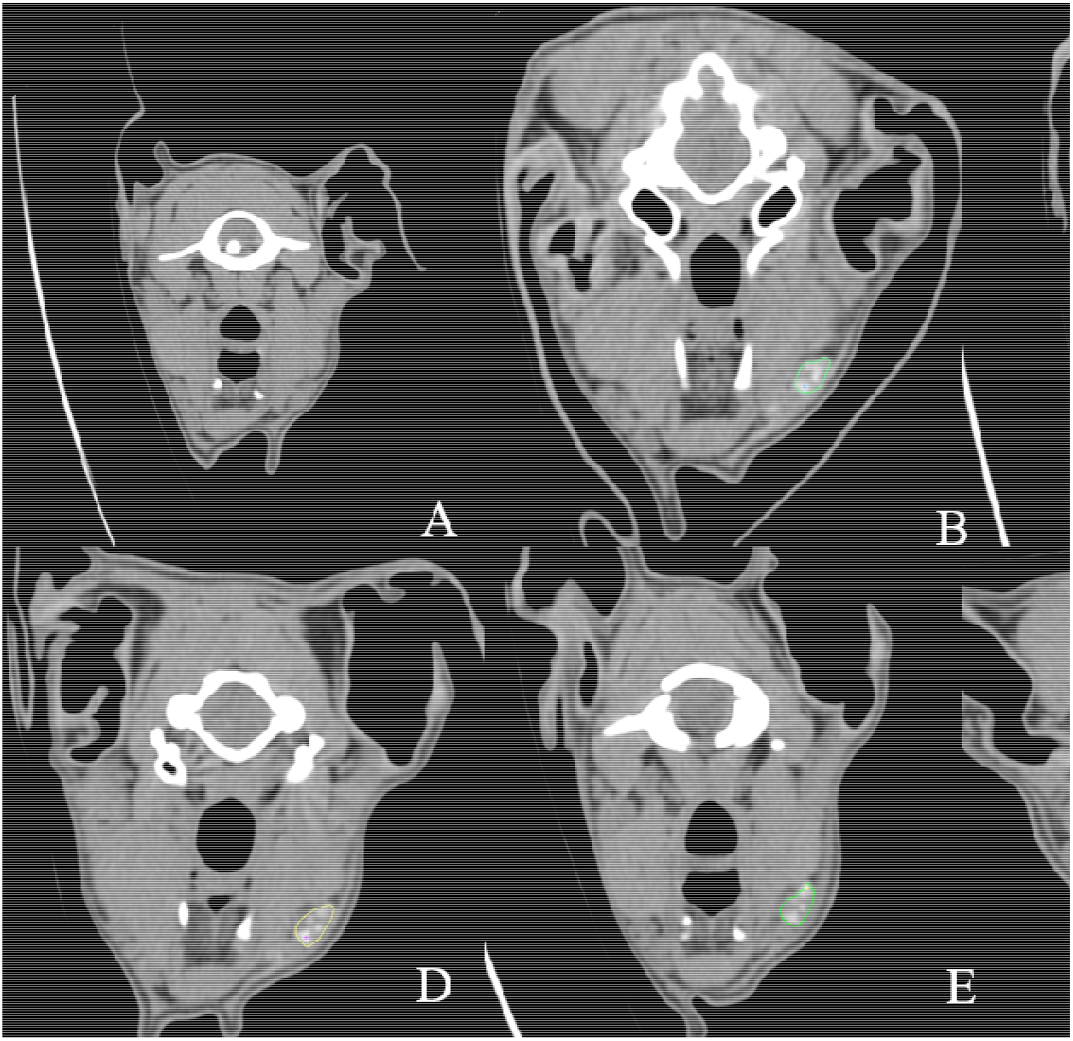
Serial cross-sectional CTL images in a transverse plane set in a soft tissue window of the same dog compared to pre-injection (A). Images B-F demonstrate contrast enhancement present within the right dorsolateral mandibular lymph node (colored outline) at day 1 post-injection (B), day 2 post-injection (C), day 4 post-injection (E), and day 10 post-injection (F).

**Figure 2.**
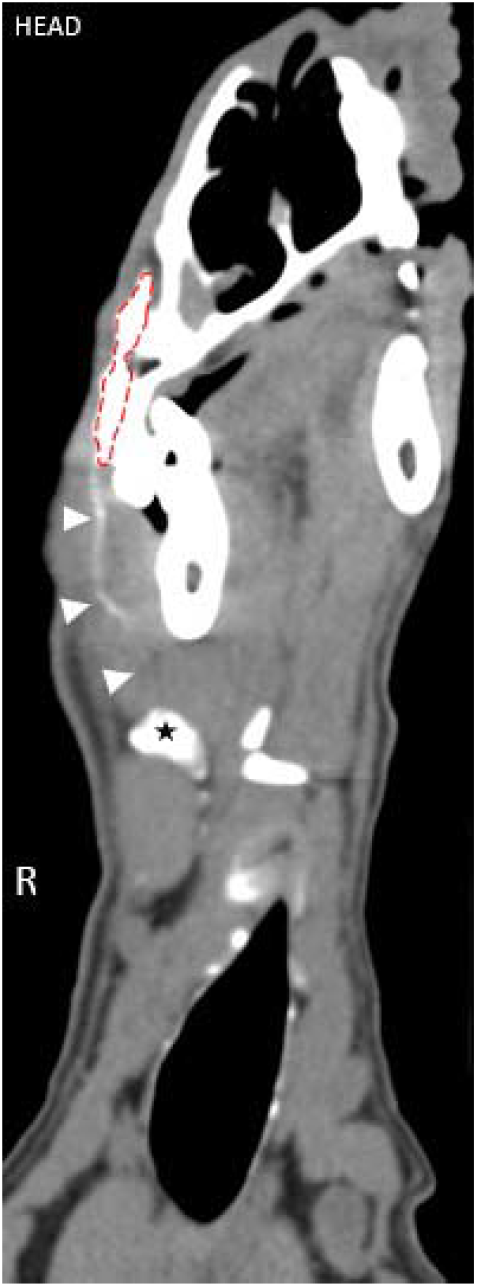
Cross-sectional CTL image in a dorsal plane set in a soft tissue window of a dog in group 2 at 1 minute post-injection demonstrating the presence of an efferent lymphatic tract (arrow heads) coursing from the injection site (red dotted outline) to the right dorsolateral mandibular lymph node (star). See Supplementary Video 1 for corresponding percutaneous fluorescence using NIRF.

The overall mean ROI volumes of the injection site were 0.94cm^3^ (group 1) and 1.20cm^3^ (group 2). There was a significant difference in mean ROI volume at all time points when compared to baseline (p<0.0001); however, there was no significant difference between groups. The overall mean ROI volume of the cervical lymph nodes was 0.27cm^3^ (group 1) and 0.19cm^3^ (group 2). There was no significant difference between groups at all time points; however, when mean ROI volume of the lymph node was analyzed within each group, a significant difference was identified within group 2 starting at day 1 p.i compared to baseline (p<0.05); however, no significant difference was found within group 1.

The overall peak mean contrast enhancement at the injection site was 910.619 HU (range 697.94-1188.10 HU) at 1 min p.i. and the peak mean contrast enhancement of the SLNs was 73.98 HU (range 63.45-86.27 HU) at 48 hours p.i. (Figure 3). Based on ANOVA for repeated measures analysis, there was a significant difference in overall mean contrast enhancement at the injection site and SLNs when compared to baseline. The mean SLN contrast enhancement within group 1 was 49.96 HU (range 40.15-62.18 HU) and in group 2 was 57.13 (42.54-71.09 HU). No significant difference in overall mean peak contrast-enhancement of the injection site and SLN was identified between groups (p=0.71) (Figure 4).

**Figure 3.**
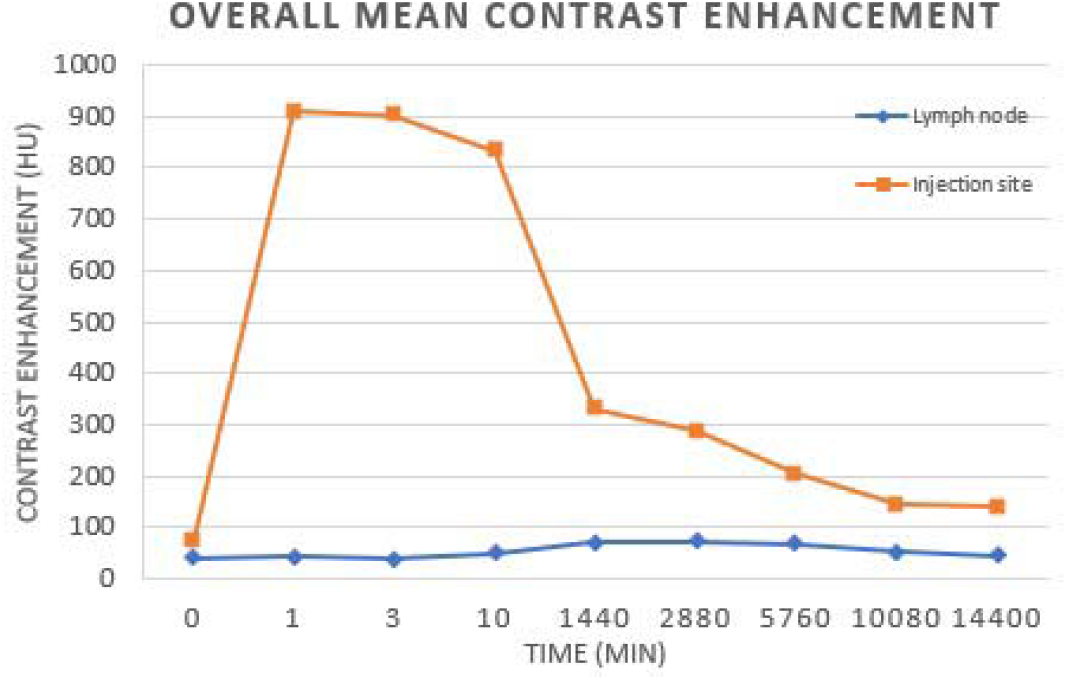
Overall mean contrast enhancement of the injection site and lymph nodes. The peak mean contrast enhancement of the lymph nodes was 73.98 HU at 2 days post-injection. There was a significant difference in SLN contrast enhancement at each time point compared to baseline (p<0.01). The peak mean contrast enhancement at the injection site was 910.86 HU at 1 minute post-injection. There was a significant difference in contrast enhancement at the injection site at each time point compared to baseline (p<0.001).

**Figure 4.**
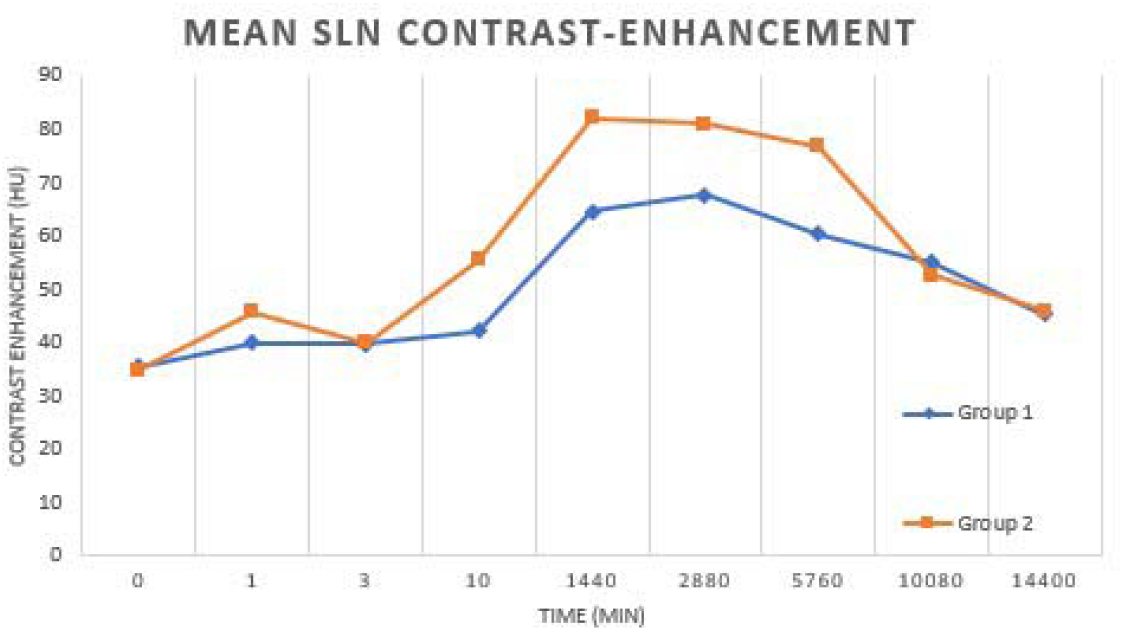
The overall peak mean contrast enhancement of the SLNs was 73.98 HU at 2 days post-injection. There was a significant difference in SLN contrast enhancement at each time point compared to baseline (p<0.01). There was no significant difference identified between groups (p = 0.71)

### NIRF

Fluorescence was detected at the injection site in all dogs at all time points and was still present at day 10 p.i (Figure 5). Using ImageJ analysis, a corrected total fluorescence measured in pixels was recorded. There was no significant difference in the overall corrected total fluorescence at each time point.

**Figure 5.**
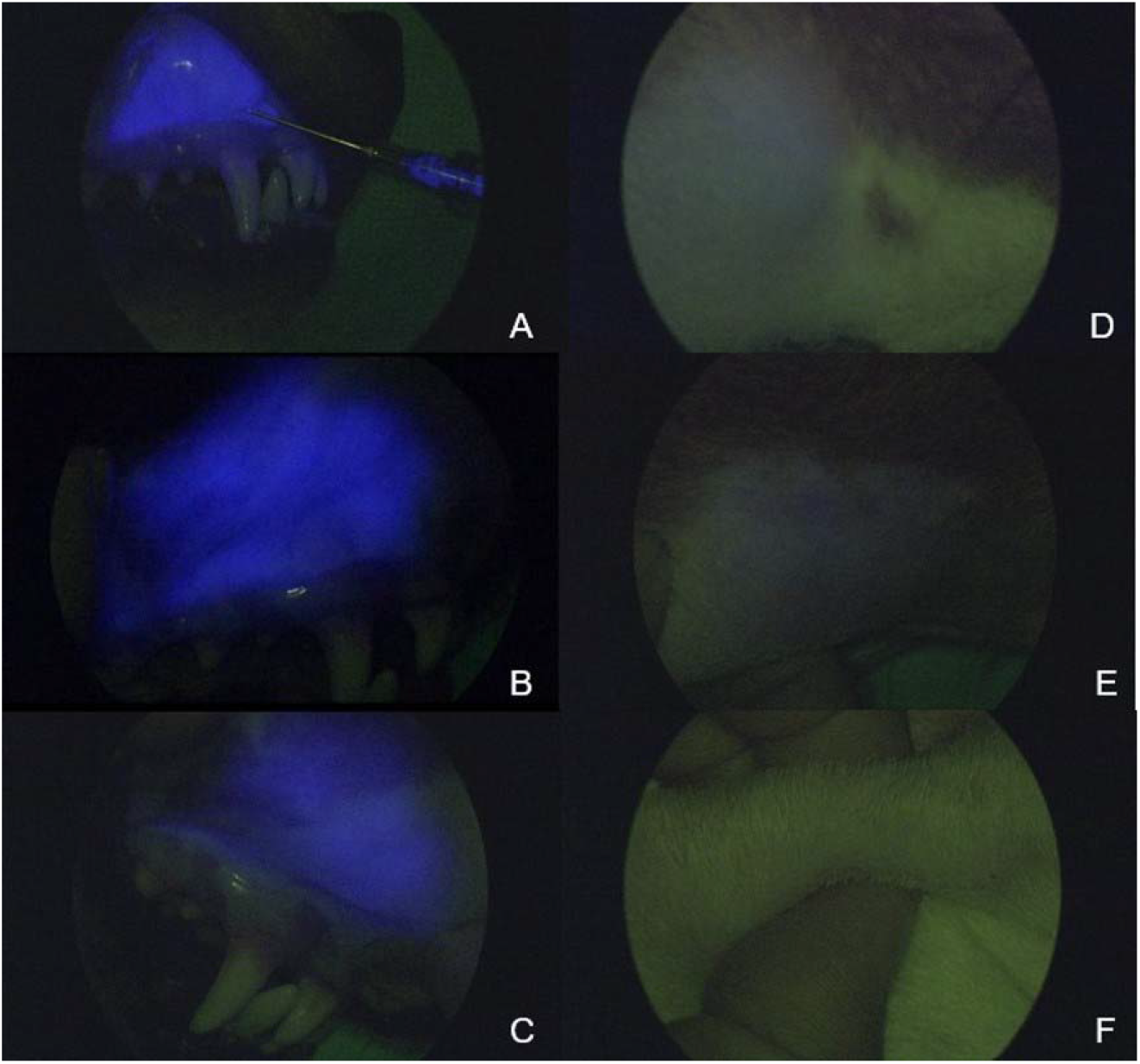
Serial near infrared fluorescence images using a NIR exoscope. Fluorescence appears as blue based on the exoscope algorithm and is present at the injection site (A-C) during injection administration (A), 4 days post-injection (B), and 10 days p.i (C). Images D-F demonstrate percutaneous fluorescence present at the right mandibular lymphocentrum at day 1 (D) and day 4 p.i (E). There is no longer percutaneous fluorescence detected within the right mandibular lymphocentrum at day 10 p.i (F).

Percutaneous fluorescence of the right mandibular lymphocentrum was also detected in all dogs (Figure 5). The mean time to detection of SLN was 6.16 ± 0.72 minutes with an overall median fluorescence intensity of 1+ in both groups. Based on a general linear mixed model, there was a significant difference between degree of fluorescence and time (p=0.0061) with degree of fluorescence significantly reduced at days 7 and 10 p.i. compared to the day of injection (p=0.0027). There was no significant difference in fluorescence intensity between groups; however, subjectively, the fluorescence intensity appeared greater in group 2 compared to group 1 immediately following injection. In half of the dogs, fluorescence was still present within the right mandibular lymphocentrum by day 7 p.i. By day 10 p.i., no dogs had detectable percutaneous fluorescence. The proportion of fluorescent lymph nodes was 96% up to day 4 p.i. compared to after 4 days (p=0.0015).

Percutaneous lymphatic tracts were identified in group 2 dogs only immediately p.i., but not in group 1 dogs; however, this difference was not found to be significant (p=0.13) (Supplementary Video 1). Percutaneous lymphatic tracts were present in 45% of cases up to 4 days p.i. compared to after 4 days (6%), which was significant (p=0.04).

### Adverse Events

There were no changes in physical examination parameters between baseline and day 7 p.i. for both groups. Based on complete blood count and serum biochemistry results, there were no clinically significant abnormalities detected between day 7 and baseline values. Evaluation of the local injection site revealed mild swelling and erythema immediately following injection, which resolved within 24 hours.

## Discussion

Our study demonstrates that the submucosal administration of CF800 within the oral cavity of healthy dogs was feasible and effective for image-guided surgery and allowed for the use of both preoperative CTL and NIRF modalities for SLN mapping.

Sentinel lymph nodes were identified in all dogs using CTL and NIRF. The combination of preoperative CTL and intraoperative NIRF for SLN mapping have been suggested to improve the accuracy for SLN detection in oncologic staging in human medicine. In both human and veterinary medicine, the combination of CTL with NIRF^15,27^ or lymphoscintigraphy and methylene blue dye^28^, the SLN detection rate improved to 100%. Preoperative CTL provides rapid imaging of contrast enhanced lymphatic tracts and SLNs, as well as, high spatial resolution and anatomic detail, which is beneficial for surgical planning.^14–16,27–29^ As the SLN may not always be the ipsilateral regional lymph node, having advanced knowledge of the targeted lymph node will be important to ensure accurate sampling. When the SLN is far outside of the standard sampling basin, this information can alter preoperative preparation and surgical approach, making advanced knowledge critical.^29^ Preoperative CTL should then be paired with an intraoperative imaging modality in order to guide the surgeon to the correct SLN for biopsy as there can often be several within a single lymphocentrum, as seen with the mandibular lymph nodes.

The median time to detection of contrast-enhanced SLNs based on CTL was 5.5 minutes, which is slightly prolonged compared to previously reported studies. Canine studies investigating indirect CTL using water soluble iodinated contrast identified SLNs within 1-3 minutes following local injection despite variations in contrast volume and injection technique.^28–30^ The minor difference reported in our study could be associated with the liposomal nature of the novel contrast agent affecting transit time. In one case, the SLNs could not be identified until the 1 day p.i. This delayed drainage but later visualization highlights a potential strength of CF-800. Since the contrast is retained in the SLN for a prolonged period, if a SLN is not identified on the immediate scans, there is still the opportunity to have a successful study on a follow-up scan without requiring re-injection. The overall mean peak contrast enhancement within the SLN was 73.98 HU which is much lower compared to previously reported studies. A potential factor contributing to a lower HU may be associated with the co-encapsulation of iohexol and ICG. These contrast agents may interfere with each other resulting in decreased contrast-enhancement or fluorescence intensity, respectively.^20^ Studies that evaluated iohexol alone for CTL of the head and neck reported HU ranging between 273.1-375.5 HU.^14,29^ Despite this marked difference in mean HU between studies, contrast-enhancement could easily be detected visually within the SLN in our study and a significant difference in contrast enhancement p.i. compared to pre-injection was identified.

The mean peak contrast enhancement of the SLN was identified at 2 days post-injection and was retained within the SLN for at least 7 days. This suggests that CTL can be performed up to 7 days p.i., but may be best detected at 2 days p.i. This finding is similar with other studies, in which CF800 was evaluated.^31,32^ In these studies, CF800 was injected intravenously in a rabbit model and was retained within the primary tumor up to 8 days p.i^31^ and 10 days p.i.^32^ Second tier lymph nodes were identified as early as 1 day post-injection, so the potential for second tier drainage must be considered if there is a prolonged period between injection and the study.

In our study, dogs that received a 2 mL volume of CF800 (group 2) had detectable contrast-enhanced (2/3) and percutaneous fluorescent (3/3) lymphatic tracts compared to dogs that received a 1 mL volume (group 1). This is an important finding as contrast uptake within the lymphatic vessels can be traced from the primary tumor to the SLN. In one study, the specific mandibular LN within the mandibular lymphatic basin could be determined from CTL based on the draining lymphatic vessels.^28^ Visualization of the lymphatic system has allowed for the identification of multiple SLNs, contralateral SLNs, and SLNs that are located away from the primary tumor^14,29^ suggesting that the closest anatomical draining node may not always be sentinel. In addition, the SLN may be differentiated from second- and third-tier lymph nodes ensuring that the correct node is identified.

The mean ROI volume of the SLNs was significantly different in group 2 dogs at days 1-7 when compared to baseline, which may suggest that contrast accumulates within the SLN. Based on our results, the authors recommend using 2 mL of CF800 for local administration to enhance contrast uptake within draining lymphatic vessels and SLN. The use of this increased volume improved visualization but there was no increase of second-tier SLN or evidence of increased morbidity associated with the increased volume.

The mean time to fluorescence within the SLN was 6.3 minutes, which is similar to a previous study.^25^ In our study, retention of fluorescence was demonstrated to be present in the SLN up to 4 days p.i.. Since no studies have evaluated the expected time of retention of ICG alone, beyond 2 hours it is difficult to directly compare retention times between ICG alone and CF-800 but it is anticipated that retention of CF-800 is significantly prolonged compared to ICG.^33^ Indocyanine green has a low molecular weight of 776 Daltons, which may contribute to its rapid transit time within lymphatic vessels and short retention time within lymph nodes.^22^ This may cause “spillover” of contrast into second- and third-tier lymph nodes, which may lead to the misidentification of these nodes as sentinel. As a result, the recommendation is for ICG to be administered at the time of surgery to allow for real-time visual guidance and extirpation of the SLNs.^34^ This fast transit precludes the ability to perform preoperative imaging and planning under the same injection. Human studies have investigated combining ICG with human serum albumin to improve retention time and increased fluorescence intensity within SLNs; however, no significant differences were identified when compared to ICG alone.^21^

Liposomes have been utilized as carriers for various drugs and contrast agents.^35^ Their use in optical imaging with ICG has demonstrated improved uptake and retention within the lymphatic system, as well as enhanced fluorescence intensity.^35,36^ The increased retention time demonstrated in our study could be due to the addition of a liposomal carrier. In a preclinical animal model study, CF800 was found to retain within metastatic lymph nodes up to 4 days p.i following intravenous injection.^23^ These findings, in conjunction with the results of our study, demonstrate that NIRF can be performed several days following intravenous or local administration of CF800 and stay retained within the SLN for at least that period of time.

Liposomes have been suggested to initiate the innate immunity resulting in hypersensitivity reactions, which have been reported in dogs with IV administration of a liposome based carrier.^37,38^ This reaction can manifest as changes to the cardiovascular system and skin, which are typically mild and transient. Rarely, they can result in severe cardiopulmonary effects.^37^ The dogs in this study were monitored for changes in cardiovascular parameters during administration of CF800 and the injection site was monitored daily for any local changes. No systemic adverse effects were encountered. The injection site was noted to be mildly erythemic at the time of injection in all dogs, but was self-limiting and resolved within 24 hours.

We have demonstrated that CF800 achieves a high SLN detection rate utilizing both preoperative CTL and intraoperative NIRF with a prolonged retention time. The application of this novel contrast agent would be advantageous in a clinical setting for several reasons. Utilizing a dual-modality contrast agent will allow for the administration of a single injection at the target of interest, thus, reducing/minimizing variations in injection technique when multimodal SLN mapping techniques are performed. The prolonged retention time will allow for these imaging procedures to be performed on separate days yet still allow for accurate and consistent SLN identification between modalities.

There are potential limitations associated with this study including the inherent nature of a preclinical experimental study. Young healthy dogs were used in our study, thus, results may differ in clinical patients with primary oral tumors, in which lymphatic drainage may be disrupted due to the presence of neoplastic cells.^36^ Cervical lymphadenectomy was not performed, thus, evaluation for fluorescence uptake within deeper cervical lymph nodes (i.e. medial retropharyngeal node) was not assessed and it is likely that as a result we underestimated the fluorescence retention time. Lastly, as CF800 has not yet been investigated in dogs, a validated dose or volume has not yet been established.

## Conclusions

The local administration CF800 has been demonstrated to be a simple and feasible technique for utilizing preoperative CTL and intraoperative NIRF for SLN mapping in the oral cavity of healthy dogs with a single injection. Retention of the contrast agent within the SLN was present for at least 4 days for percutaneous fluorescence and 7 days for CT enhancement. Later-tier SLNs occurred in 4/6 dogs but did not occur until 24 hours following injection. In one dog, a 3rd-tier SLN was identified 10 days following injection. CF800 was found to be safe to use with minimal to no local adverse reactions. The application of this novel contrast agent may facilitate preoperative and intraoperative SLN mapping in dogs with tumors of the head and neck; however, further investigation in clinical patients is warranted.

## Acknowledgements

The authors would like to thank Gabrielle Monteith, Research Statistician, Department of Clinical Studies, Ontario Veterinary College, University of Guelph, for her contributions to statistical analysis and interpretation.

The authors declare no conflict of interest related to this report.

## Supplementary Video 1

This video clip was recorded using a NIR exoscope from the same dog as in Figure 2. The dog is positioned in left lateral recumbency with the head towards the right of the screen. The video clip demonstrates the presence of percutaneous fluorescence at the injection site (submucosa oral cavity) followed by percutaneous fluorescent lymphatic tracts coursing along the right ventrolateral neck from the injection site to the right mandibular lymph node.

